# Testing the exteroceptive function of nociception: the role of visual experience in shaping the spatial representations of nociceptive inputs

**DOI:** 10.1101/536367

**Authors:** Camille Vanderclausen, Marion Bourgois, Anne De Volder, Valéry Legrain

## Abstract

Adequately localizing pain is crucial to protect the body against physical damage and react to the stimulus in external space having caused such damage. Accordingly, it is hypothesized that nociceptive inputs are remapped from a somatotopic reference frame, representing the skin surface, towards a spatiotopic frame, representing the body parts in external space. This ability is thought to be developed and shaped by early visual experience. To test this hypothesis, normally sighted and early blind participants performed temporal order judgment tasks during which they judged which of two nociceptive stimuli applied on each hand’s dorsum was perceived as first delivered. Crucially, tasks were performed with the hands either in an uncrossed posture or crossed over body midline. While early blinds were not affected by the posture, performances of the normally sighted participants decreased in the crossed condition relative to the uncrossed condition. This indicates that nociceptive stimuli were automatically remapped into a spatiotopic representation that interfered with somatotopy in normally sighted individuals, whereas early blinds seemed to mostly rely on a somatotopic representation to localize nociceptive inputs. Accordingly, the plasticity of the nociceptive system would not purely depend on bodily experiences but also on crossmodal interactions between nociception and vision during early sensory experience.

## 1. Introduction

Pain is as an unpleasant sensory and emotional experience associated with actual or potential tissue damage (IASP, 1994). It usually results from the activation of nociceptors, sensory receptors characterized by high activation thresholds, i.e. by the capacity to respond – at least under normal conditions – to stimuli of high intensity and potentially noxious (Belmonte & Viana, 2008). Pain has therefore an interoceptive function of warning the brain about the occurrence of sensory events having the potential to damage the body (Craig, 2003). Among other functions of pain, its localization on the skin or in the viscera is of primary importance because it helps to identify which part of the body is being damaged. Conversely to the classical view (e.g. Kandel, Schwartz, Jessell, Siegelbaum, & Hudspeth, 2013), nociceptive inputs can provide detailed and accurate spatial information, suggesting finely-tuned mapping systems for pain (Moore & Schady, 1995). Surprisingly, studies having investigated the mapping organization of nociceptive inputs in the brain (Andersson, 1997; Baumgartner et al., 2010; Bingel et al., 2004; Henderson, Gandevia, & Macefield, 2007; Mancini, Haggard, Iannetti, Longo, & Sereno, 2012) only focused on the somatotopic organization characterized by anatomical representations of the body surface based on the ordered projection of the receptor fields to spatially segregated subgroups of neurons (Penfield & Rasmussen, 1950). However, it has been repeatedly shown that innocuous tactile inputs can be recoded according to spatiotopic representations, i.e. representations that use external space as reference frame, taking the relative position of the limb on which a given stimulus is applied into account (Azanon & Soto-Faraco, 2008; Graziano, Hu, & Gross, 1997; Heed & Azañon, 2014; Iwamura, Tanaka, Sakamoto, & Hikosaka, 1993; Shore, Spry, & Spence, 2002; Smania & Aglioti, 1995; Yamamoto & Kitazawa, 2001). While somatotopic maps allow coding the position of contacts on the skin surface, spatiotopic maps provide an appropriate readout for the brain allowing to identify the object in external space that is in contact with the body, and therefore planning an adequate spatially guided action towards that object (Brozzoli, Ehrsson, & Farne, 2014). Such complex ability to represent somatic information appears even more crucial for nociceptive and painful stimuli since it allows to detect and appropriately react against noxious stimuli that threathen the physical integrity of the body (Legrain & Torta, 2015). Demonstrating the brain’s ability to map nociceptive inputs according to spatiotopic representations would provide evidence for the exteroceptive function of nociception (Haggard, Iannetti, & Longo, 2013), whose role would be to optimize the monitoring of space around the body and react to potential danger (Legrain, 2017).

Spatiotopic mapping of touch has been recurrently evidenced using temporal order judgment (TOJ) tasks during which participants judge the order of apparition of two successive tactile stimuli, one applied to each hand, and separated by different temporal delays (Heed & Azañon, 2014). It is noteworthy that TOJ tasks are performed with the hands either in a normal uncrossed posture or crossed over the midsagittal plane of the body. Participants’ judgements are typically less accurate when their hands are crossed, and such an effect is accounted by the fact that the somatotopic representation (“*Which hand is stimulated?*”) mismatches the spatiotopic representation (“*Where is the stimulated hand?*”) (Shore et al., 2002; Yamamoto & Kitazawa, 2001). This indicates that, when judging the position of a tactile stimulation on the body, its position is automatically recoded according to spatiotopic frames of references (Azanon & Soto-Faraco, 2008; Heed & Azañon, 2014). Importantly, it has been suggested that spatiotopic representations of touch are not innate but would rather develop during infancy (Azanon, Camacho, Morales, & Longo, 2017; Pagel, Heed, & Röder, 2009). Accordingly, hand posture does not affect the performance of people with early and complete visual deprivation, suggesting that the ability to remap touch according to external frame of reference is, at least partially, shaped by early visual experiences (Crollen, Albouy, Lepore, & Collignon, 2017; Röder, Rösler, & Spence, 2004).

The first aim of the present experiments was to test the hypothesis according to which nociceptive inputs are automatically coded according to spatiotopic reference frames. Normally sighted participants performed temporal order judgment tasks on thermal stimuli specifically and selectively activating skin nociceptors. Stimuli were applied on each hand dorsum and tasks were performed with the hands either uncrossed or crossed. We expected a decrease of performance in the crossed posture as compared to the uncrossed posture. The second aim was to test the hypothesis of the role of early visual experience in the development of the spatiotopic representation of nociception. We therefore compared the ability of early blind participants and matched blindfolded sighted controls in discriminating the temporal order of nociceptive stimuli. Considering that touch and nociception share the same spatial representation (Legrain & Torta, 2015), we would expect early blind participants to be unaffected by hand posture, indicating that their judgements mostly rely on somatotopic representations of nociception. However, due to the higher relevance of nociception in terms of survival and the underlying fundamental role of its spatial representation, we could also expect that early blind participants would be affected by the conflict between somato- and spatiotopic representations during crossed hand posture. In this line, the spatial mapping of nociceptive stimuli would be less dependent on external factors, such as early visual experience, than that of tactile inputs.

## 2. Materials and methods

### 2.1. Participants

Thirteen healthy volunteers took part to Experiment 1. One participant was excluded because he could not achieve task requirements properly (see Procedure). The mean age of the 12 remaining participants (7 women) was 25.00 ± 2.63. Nine participants were right-handed and three of them were left-handed according to the Flinders Handedness Survey (Nicholls, Thomas, Loetscher, & Grimshaw, 2013). The participants had normal to corrected-to-normal vision. They did not report any prior history of severe neurological, psychiatric or chronic pain disorders. They did not suffer from a traumatic injury of the upper limbs within the six months nor any cutaneous lesion on the hands’ dorsum. The regular use of psychotropic drugs or the intake of analgesic drugs (e.g. NSAIDs and paracetamol) within the twelve hours preceding the experiment were also considered as exclusion criteria.

Thirteen early blind as well as sixteen normally sighted individuals participated to Experiment 2. One of the early blind and three of the normally sighted participants were excluded from the data set because they could not achieve the task requirements properly (see Procedure). Among the 12 remaining early blind participants (7 women, 36.67 ± 10.98 years of mean age), there were 9 right-handed, 1 ambidextrous and 2 left-handed and participants (Nicholls et al., 2013). The 12 remaining sighted participants (36.08 ± 10.73 years) were matched individually to the early blind participants in terms of age, sex, and level of education. The inclusion and exclusion criteria for sighted participants were similar as for Experiment 1. In addition, the early blind participants were recruited according to blindness attributed to peripheral deficits without additional neurological problem (see Table 1 for a detailed description of blind participants). They were all considered as totally blind since birth. One of them, participant EB5, because of a genetic disease, became totally blind at 3 months of life, but was still considered as early blind since his visual acuity was very poor in the first months of life. The participant EB9 of the blind group suffered from epilepsy and was under medication for that reason (Levetiracetam 1750 mg and Oxcarbazepine 1500 mg daily). The epileptic focus was located in the left temporo-occipital border of the cortical brain. Nevertheless, this participant was not excluded from the study since there was no evidence of cognitive impairment or somatosensory perception deficit. The participant EB12 suffered from attention deficit and hyperactivity disorder (ADHD) but he was free of medication (Rilatine) at the moment of the experiment.

**Table 1:**
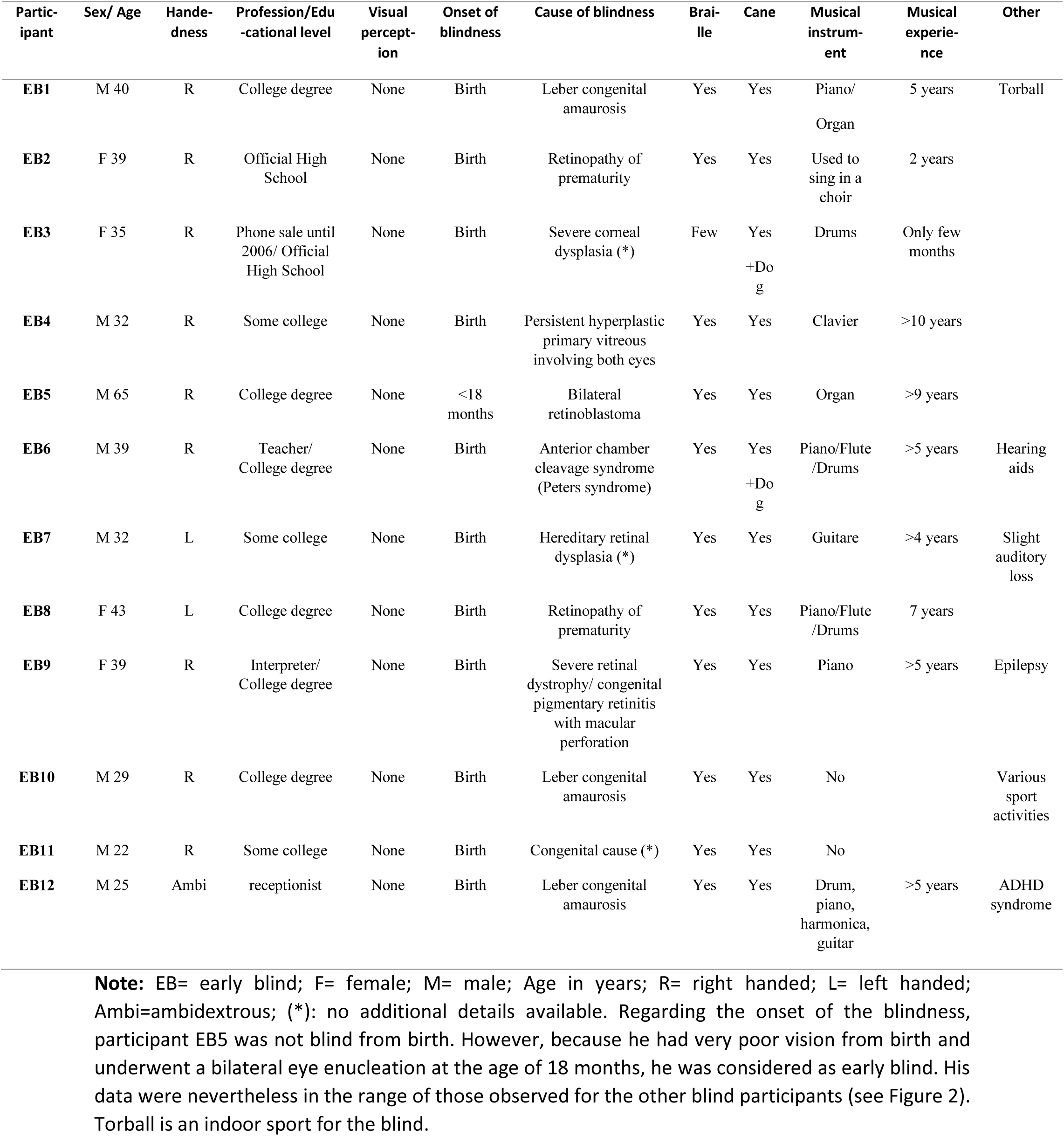
Profile of the blind subjects.

All experimental procedures were approved by the local ethics committee and conformed to the Declaration of Helsinki. Written informed consents were obtained for all participants before starting the study.

### 2.2. Stimuli and apparatus

Nociceptive stimulations consisted of radiant heat stimuli delivered onto the skin of the hands’ dorsa by means of two identical infrared CO_2_ laser stimulators (wavelength 10.6 *µ*m; Laser Stimulation Device, SIFEC, Ferrières, Belgium). This technique is known to selectively and specifically activate thermo-sensitive cutaneous nociceptors (Plaghki & Mouraux, 2005). The power of the output stimulation was regulated using a feedback control based on an online measurement of the skin temperature at the site of stimulation by means of a radiometer whose field of view was collinear with the laser beam. This allows defining specific skin temperature profiles (see Churyukanov, Plaghki, Legrain, & Mouraux, 2012). The laser beams were conducted through 10-m optical fibers. Each fiber ended with a head containing the optics used to collimate the laser beam to 6 mm diameter at the target site. Each laser head was hold upon each participant’s hand by means of articulated arms attached to a camera tripod system (Manfrotto, Cassola, Italy). Each laser head was fixed into a clamp attached to a 3-way head allowing displacements of the target site of the laser beam perpendicularly oriented to the hand dorsum by means of several sliders in all directions. During threshold measurements and during the experiments, laser beams were displaced after each stimulus using the sliders. Stimulus duration was 100 ms. Stimuli were composed of a 10-ms heating ramp dedicated to reach the target temperature, followed by a 90-ms plateau during which the skin temperature was maintained at the target temperature. Heating was then stopped. The target temperature was determined for each participant’s hand according to individual activation threshold of nociceptive thinly myelinated Aδ fibers. Thresholds were estimated by means of an adaptive staircase procedure using reaction times (RTs) to discriminate detections triggered by Aδ fiber inputs (RT < 650 ms) from detections triggered by C-fiber inputs (RT ≥ 650 ms) (Churyukanov et al., 2012). Indeed, Aδ fibers have faster nerve conduction velocity than unmyelinated C fibers (~10 m/s vs. ~1 m/s), while Aδ-fibers are known to have higher thermal activation thresholds than C-fibers (Plaghki & Mouraux, 2005). Concretely, the participants were asked to press a button with the non-stimulated hand as soon as they felt something on the stimulated hand. Any RT equal or superior to 650 ms leaded to a temperature increase of 1°C for the next stimulus. On the contrary, any RT inferior to 650 ms leaded to a decrease in temperature of 1°C. The procedure started at 46°C and lasted until four reversals were encountered. The mean value of the four temperatures that leaded to a reversal was considered as the threshold. For the stimuli used during the experimental phase, 5°C were added to that value, and if necessary, the temperature was slightly adapted for each hand and across experimental blocks so that stimuli were always perceived as equally intense between the two hands. View of the hands was prevented in sighted participants during threshold estimation. Stimuli at such temperature values were perceived as pricking and elicited a slightly painful sensation. Before starting and during the experiment, elicited sensations were tracked using a list of words to be chosen (not perceived, light touch, tingling, pricking, warm, burning). Subjective intensity was measured for each hand using a numerical rating scale (NRS) from 0 (no sensation) to 10 (strongest sensation imaginable). This was made to ensure that stimuli were still perceived as pricking and equally intense between the two hands. Stimuli temperatures were then adapted if necessary.

### 2.3. Procedure

The procedure was exactly the same for Experiments 1 and 2. The participants were sitting on a chair in front of a table. Their hands rested on the table palms down, with a distance of ~30 cm between the two index finger tips. Participants’ head was placed in a chin-rest in order to minimize head movement during the experiment. The participants were asked to perform the task with their hands either uncrossed or crossed over the body midline. Noises from experimental devices were covered by means of a white noise played through earphones during the whole experiment. The sighted participants were blindfolded with an eye mask.

Participants were presented with four blocks of 40 trials each. During two blocks, the task was performed with the hands in an uncrossed posture, whereas in the two other blocks it was performed with the hands crossed over the sagittal midline of their body (the order of these two conditions was counterbalanced across participants). Each trial consisted in pairs of nociceptive stimuli, one applied on each hand, separated in time by 24 possible stimulus onset asynchronies (SOA): ±10, ±15, ±30, ±45, ±60, ±75, ±90, ±150, ±200, ±400, ±500, ±600 ms. Negative values indicated that the first stimulus of the pair was applied on the left hand, and the second one on the right hand. Positive values indicated that the right hand was stimulated first. Within each block and for each trial, the presented SOA was determined online based on participants’ performance on all previous trials according to the PSI method (Kingdom & Prins, 2010; Kontsevich & Tyler, 1999). Based on a Bayesian framework, this adaptive procedure estimates the posterior distribution of the parameters of interest and minimizes its expected entropy (i.e. uncertainty) trial by trial, considering all the previous trials. In other words, at each trial, the algorithm infers which condition (i.e. SOA) is the most informative to estimate the distribution of the parameters of interest. This method thus allowed us to estimate the parameters of interest without probing extensively all the possible SOAs, which would be time consuming (Filbrich, Alamia, Burns, & Legrain, 2017). After each trial, the participants had to report verbally which one of the two stimuli they perceived as being presented first in half of the experimental blocks (‘which is first’), and which one they perceived as being presented second in the other half (‘which is second’), by saying ‘left’ or ‘right’ aloud. The order of these instructions was randomized within each posture condition. The aim of using two response modalities was to prevent the data from being influenced by a response bias (Filbrich, Torta, Vanderclausen, Azanon, & Legrain, 2016; Spence & Parise, 2010). Participants’ responses were encoded by the experimenter by using a keyboard, and the next trial started 2000 ms later. The time interval between two trials varied from 5 to 10 seconds. Laser beams were displaced after each trial between the heating offset and encoded experimenter’s response, which triggered the next trial. The task was unspeeded but the participants were told to be as accurate as possible. They did not receive any feedback on their performance in the task.

Experiments were preceded by a practice session of 4 blocks of 5 trials each, one block per hand posture (i.e. uncrossed vs. crossed) and response condition (i.e. ‘which is first’ vs. ‘which is second’), but only with two SOAs among the largest SOA (±150 and ±200 ms). During the experiment, to avoid overheating of the skin and habituation, a 10 minutes break was imposed to the participants between the blocks. One block lasted 10 to 15 minutes. The whole study, including the threshold measurement, the training session and the experiment lasted two to three hours.

### 2.4. Measures

Aδ fiber activation thresholds and stimulation intensities (corresponding to the averaged intensity used for each hand across all experimental blocks (i.e. approximately 5°C added to the Aδ fiber activation threshold) were measured in degree Celcius (°C). Regarding temporal order judgment (TOJ) performance, for all experimental conditions, the proportion of left stimuli having been reported as presented first was calculated as a function of SOA. For each participant, data were fitted online during the experimental block with the logistic function, i.e. *f(x)* = 1/(1+exp(-β(x-α)), from which the parameters of interest were computed (Filbrich, Alamia, Burns, et al., 2017). These parameter estimates corresponded to the last update computed by the logarithm during the adaptive procedure (Kontsevich & Tyler, 1999). They characterized respectively the threshold (α) and the slope (ß) of the function. In the present experiments, the parameter of interest was the ß parameter, describing the noisiness of the participants’ responses, i.e. the precision of their responses during the experiment (Kingdom & Prins, 2010). The β is classically used to derive from TOJ performances the just noticeable difference (JND) that denotes the SOA needed for the participants to correctly perceive the order of the two stimuli in a certain percentage of trials (Crollen, Albouy, et al., 2017; Röder et al., 2004; Shore et al., 2002). Although the contribution of the α parameter was out of the scope of the present study, it was taken into account in order to assess the presence of potential biases which could influence the estimation of the slope in the frame of the adaptive PSI method (Kingdom & Prins, 2010). The α is the threshold of the function and refers to the point of subjective simultaneity (PSS), defining the SOA at which the participants report the two stimuli as occurring first equally often. In other words, in the present experiments, The α corresponded to the SOA at which the proportion of trials during which the stimuli applied to the left hand were reported as presented first reaches 0.5. Since the PSI method is based on a Bayesian approach, a prior probability distribution needs to be postulated, based on previous knowledge regarding the values of the parameter of interest (Filbrich, Alamia, Burns, et al., 2017; Kingdom & Prins, 2010). In the present experiments, we used a prior distribution of 0 ±20 for the α parameter and 0.06 ±0.6 for the ß parameter (Filbrich, Alamia, Burns, et al., 2017). Because the adaptive PSI method was used, a third parameter was computed and corresponded to the mode of the presented SOAs, i.e the value of the SOA that was the most frequently presented to each participant during each adaptation procedure. This measure can be considered as indexing the participant’s adaptation during the task, with the idea that smaller is the mode, better is the performance, and larger it is, worse is the performance because this indicates that, overall, the participant needed higher SOAs to be able to discriminate between the two stimuli.

### 2.5. Analyses

Data were excluded from further statistical analyses if the slope of the psychometric function could not be reliably estimated during the 40 trials within one condition. Since different responses were used to minimize potential response biases, the data from the two response conditions (‘which is first’ and ‘which is second’) were merged (i.e. averaged).

Analyses were first performed on the Aδ fiber activation threshold and stimulation intensity values, to ensure that no difference was found between both hands regarding these parameters that could have influenced the results in any way. In Experiment 1, comparison between activation thresholds and stimulation intensities was made using paired t-tests with the *hand* as factor (left vs. right). In Experiment 2, we used an analysis of variance (ANOVA) for repeated measures with adding the group as the second factor (sighted vs. blind).

Regarding the TOJ tasks, in order to examine the presence of potential perceptual biases towards one of the two sides of space, one-sample t-tests were performed to compare the PSS values to 0 for each condition of the posture factor (uncrossed vs. crossed) and for each of the three groups separately. Next, PSS, slope and mode values were compared using an ANOVA for repeated measures with the *posture* (uncrossed vs. crossed) as a within-participant factor in Experiment 1, and using the *posture* and the *group* (early blind vs. normally sighted) respectively as within- and between-participant factors in Experiment 2. Greenhouse-Geisser corrections and contrast analyses were used if necessary. Effect sizes were measured using partial Eta squared for ANOVA and Cohen’s d for t-tests. Significance level was set at *p*≤ .05.

## 3. Results

### 3.1. Experiment 1

#### 3.1.1. Threshold and intensity values

Aδ-fiber activation threshold value obtained for the left hand (M=48.08±2.02°C) was not significantly different from the threshold for the right hand (M=47.58±2.11°C) (t(11)=0.88, *p*=.400, d=0.25). Similarly, there was no significant difference between the left (M=53.63±1.58) and the right (M=53.33±1.76) hands for the stimulation intensity values (t(11)=.77, *p*=455).

#### 3.1.2 TOJ values

The fitted psychometric curves are illustrated in Figure 1a. The one sample t-tests revealed that the PSS value in the crossed posture condition (M=-0.30±14.11ms) was not significantly different from 0 (t(11)=-0.07, *p*=.943, d=0.02). But, surprisingly, PSS was slightly but significantly different from 0 in the uncrossed posture condition (t(11)=2.37, *p*=.037, d=0.68). With a mean value of 8.86ms (SD=12.95), it indicated a slightly biased judgment toward the left hand in this posture condition.

**Figure 1.**
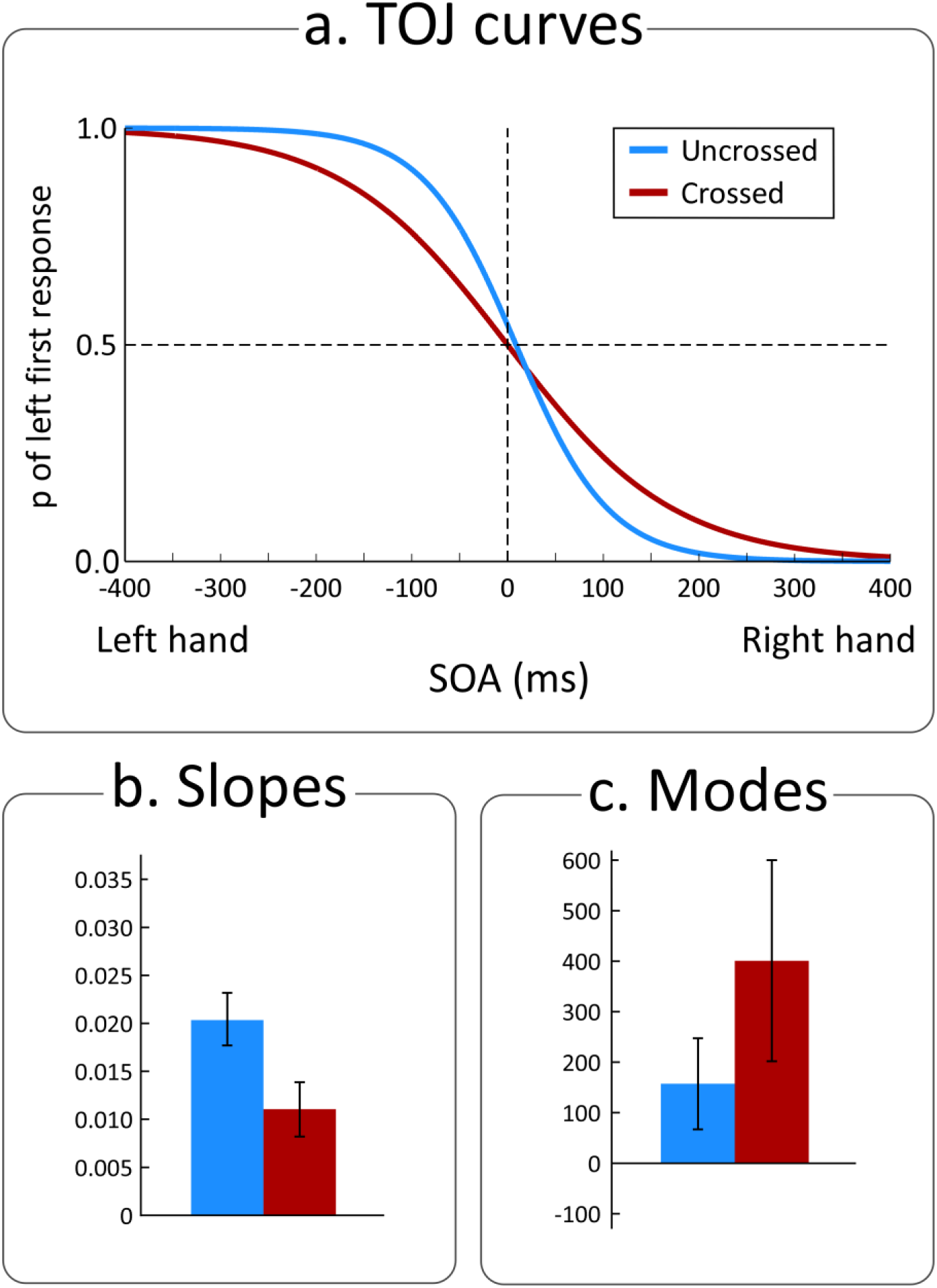
Nociceptive temporal order judgments in Experiment 1. (a) Fitted curves of the psychometric function from the data of the 12 sighted participants according to the hands posture condition. The x-axis represents the different possible SOAs. A negative value indicates that the left hand was stimulated first and a positive value indicates that the right hand was stimulated first. The y-axis represents the proportion of trials in which the nociceptive stimulus applied on the left hand was perceived as being presented first. The lines represent the fitted curves for the uncrossed posture condition (blue) and the crossed posture condition (red), respectively. (b) Mean of the slope values for each posture condition. The slope value of the crossed posture condition was significantly lower than the slope value corresponding to the uncrossed posture condition. (c) Mean of the mode values of the presented SOAs in millisecond for each posture condition. The averaged mode in the crossed posture condition was significantly higher than the mean mode value of the uncrossed posture condition. Error bars represent the 95% confidence intervals estimated for each measure according to the Cousineau’s method (Cousineau, 2005).

Comparison of the posture conditions revealed no significant effect on the PSS values (F=4.49, *p*=0.580, 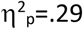), suggesting that the bias observed in the uncrossed posture condition was marginal. Regarding the slope values, the ANOVA revealed a significant effect of the posture (F(1,11)=13.13, *p*=.004, 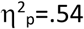), showing that the slope value was significantly lower in the crossed posture (M=0.011±0.006) than in the uncrossed posture condition (M=0.021±0.009) (Figures 1b & 2). Similarly, analysis of the mode of the presented SOAs revealed a significant effect of the posture (F(1,11)=8.82, *p*=.013, 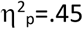). The mode of the presented SOAs was significantly larger in the crossed posture condition (M=402.50±259.13ms), as compared to the uncrossed posture condition (M=157.50±211.28ms) (Figure 1c).

### 3.2. Experiment 2

#### 3.2.1 Threshold and intensity values

Analyses of the Aδ fibers activation thresholds showed no significant effect of the hand (F(1,22)=1.12, *p*=.302, 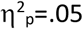), no significant effect of the group (F(1,22)=1.30,*p*=.267, 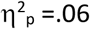), and no significant interaction between the two factors (F(1,22)=0.17, *p*=.689, 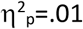). Early blind participants had a similar activation threshold (M=48.25±2.18°C) than the normally sighted participants (M=49.21±1.94°C). Similar results were obtained for the stimulation intensity, as the ANOVA revealed neither significant main effect of the hand (F(1,22)=1.71, *p*=.205, 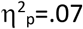) nor of the group (F(1,22)=0.94, *p*=.344, 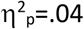), nor significant interaction between the two factors (F(1,22)<0.01, *p*=.949, 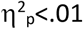).

#### 3.2.2. TOJ values

The fitted psychometric curves are illustrated for both groups in Figure 3a. The one sample t-tests performed for each group and for each posture separately showed that none of the PSS values were different from 0 (all t(11)≤-1.35, all p≥.204).

**Figure 3.**
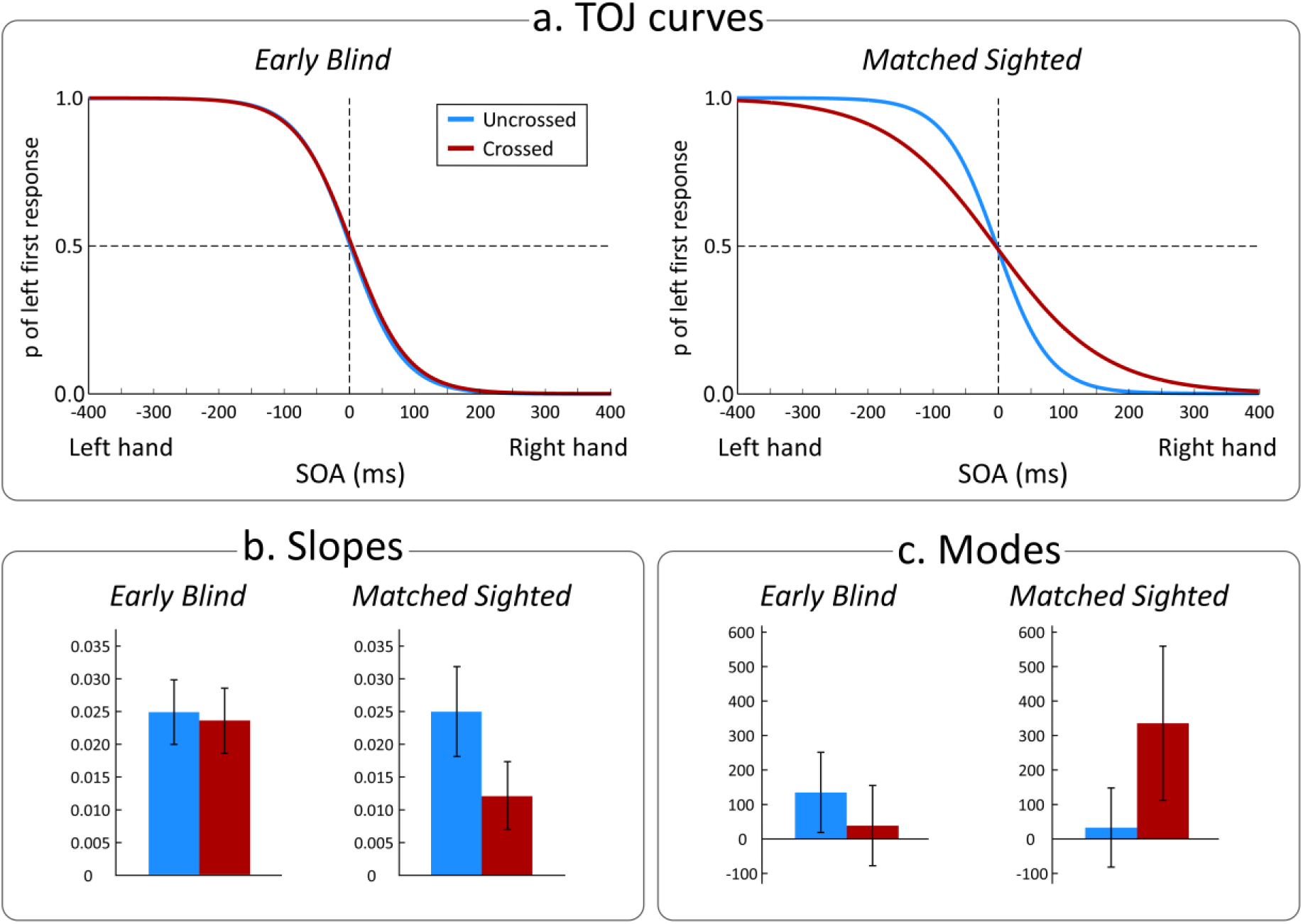
Nociceptive temporal order judgments in Experiment 2. (a) Fitted curves of the psychometric function from the data of the 12 early blind and the 12 matched sighted participants according to the hands posture condition. The x-axis represents the different possible SOAs. A negative value indicates that the left hand was stimulated first and a positive value indicates that the right hand was stimulated first. The y-axis represents the proportion of trials in which the nociceptive stimulus applied on the left hand was perceived as being presented first. The lines represent the fitted curves for the uncrossed posture condition (blue) and the crossed posture condition (red) for the early blind group and the matched sighted group, respectively. (b) Mean of the slope values for each posture condition for each group. The slope value of the crossed posture condition was significantly lower than the slope value of the uncrossed posture condition only in the sighted group. In the early blind participants, the averaged slope in the crossed and uncrossed posture conditions were not significantly different from each other. (c) Mean of the mode values of the presented SOA in millisecond for each posture condition and each group. The averaged mode in the crossed posture condition was significantly higher than the averaged mode value of the uncrossed posture condition in the sighted group but such difference was not found significant for the early blind group. Error bars represent the 95% confidence intervals estimated for each measure according to the Cousineau’s method (Cousineau, 2005).

The ANOVA performed on the PSS values revealed no significant difference between the posture conditions (F(1,22)=0.01, *p*=.922, 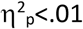), no significant difference between the groups (F(1,22)=0.77, *p*=.389, 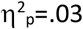) and no significant interaction between the two factors (F(1,22)=0.20, p=.657, 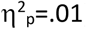). On the contrary, the ANOVA performed on the slope values revealed a significant main effect of the posture (F(1,22)=6.33, *p*=.020, 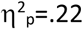) and an almost significant interaction with the group factor (F(1,22)=4.25, *p*=.051, 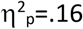). The main effect of the group was not significant (F(1,22)=3.09, *p*=.092, 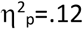). Contrast analyses revealed a significant effect of the posture in the normally sighted group (t(11)=3.76, *p*=.003, d=1.08), the slope value of the crossed condition (M=0.012±0.010) being lower than that of the uncrossed condition (M=0.025±0.009). Conversely, in the blind group, such a difference was not significant (t(11)=0.29, p=.779, d=0.08), the slope value of the crossed condition (M=0.023±0.008) being comparable to that of the uncrossed condition (M=0.024±0.013) (Figures 2 and 3b). Regarding the mode of the presented SOAs (Figure 3c), the results corroborated those of the slope values, as the ANOVA revealed an almost significant main effect of the posture (F(1,22)=4.12, *p*=.055, 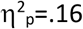) and a significant interaction with the group factor (F(1,22)=15.34 *p*=.001, 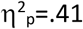), while there was no significant main effect of the group (F(1,22)=3.03, *p*=.096, 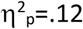). Contrast analyses revealed a significant difference between the two postures in the normally sighted group (t(11)=-3.79, *p*=.003, d=1.10), the mode of the presented SOAs being larger in the crossed posture condition (M=329.17±284.49ms) as compared to the uncrossed posture condition (M=32.92±42.13ms). The difference (crossed posture: M=36.67±45.19ms; uncrossed posture: M=130.83±222.77ms) was not significant in the early blind group (t(11)=1.53, *p*=.156, d=0.44) (Figure 3c).

**Figure 2.**
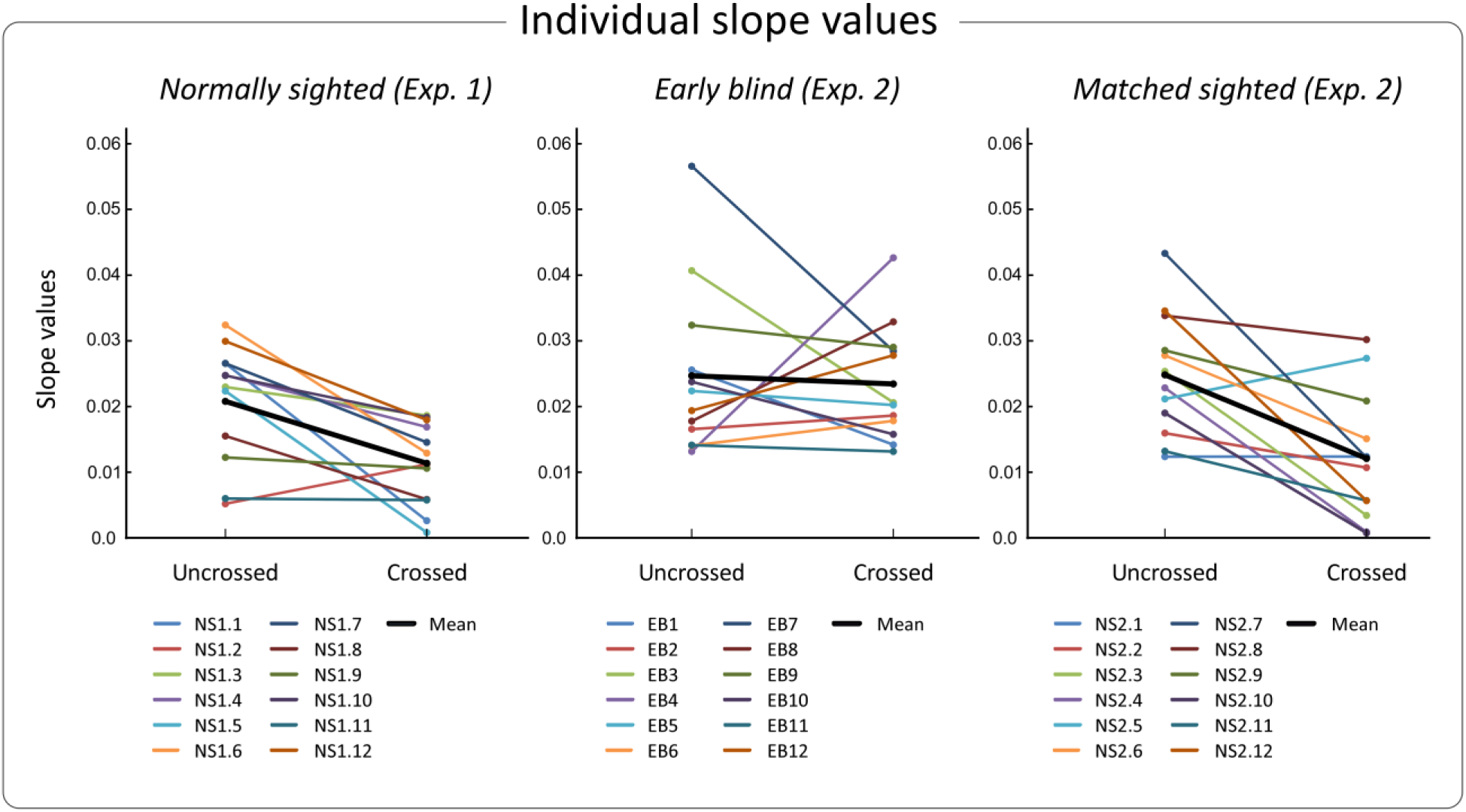
Individual data in Experiments 1 and 2. Individual slope values corresponding to the uncrossed and crossed posture conditions for each sighted observer that participated in the Experiment 1, as well as each early blind and matched sighted participants from Experiment 2. The averaged slope values for each experiment, each group and each posture condition are represented by the thick black lines. The data of participants EB5, EB9 and EB12 are in the range of the group values.

## 4. Discussion

The goals of the present study were to characterize the spatial representations of nociceptive stimuli and the role of early visual experience in shaping these representations. To this aim, two experiments were conducted by means of TOJ tasks during which participants discriminated the temporal order of two nociceptive stimuli, one applied on each hand placed in either an uncrossed or crossed posture. While early blind participants’ performance was not affected by the posture, the performance of the two groups of blindfolded sighted participants decreased in the crossed condition as compared to the uncrossed condition. As the crossed posture is aimed to generate a mismatch between the somatotopic and the spatiotopic representations of somatic inputs (Heed & Azañon, 2014), the results indicated that when normally sighted people had to localize a nociceptive stimulus on their body, its position was automatically mapped into a spatiotopic representation and interfered with the somatotopic map. Since the same pattern of results was observed with innocuous tactile stimuli (Shore et al., 2002; Yamamoto & Kitazawa, 2001), it is assumed that the perception and localization of touch and nociception, although being processed through two segregated afferent pathways, share similar spatial representations (Legrain & Torta, 2015). These results are in line with the study of Sambo et al. (2013) who showed a crossing-hands effect during TOJ tasks using intra-epidermal electrical stimulation (IES). While the selective activation of nociceptors by IES depends on a strict stimulation procedure (Mouraux, Iannetti, & Plaghki, 2010), we replicated here these results by using radiant heat stimuli delivered by means of two CO_2_ laser stimulators, a technique that allows undoubtedly specific and selective activation of skin nociceptors (Plaghki & Mouraux, 2005). More specifically, the staircase procedure based on participants’ reaction times allowed us to target nociceptors associated to Aδ fibres (Churyukanov et al., 2012). Nevertheless, the most novel result of the present study comes from the comparison between normally sighted and early blind participants. These results showed that early blind participants were unaffected by the posture of the hands as they performed the TOJ tasks similarly in the crossed and uncrossed conditions, a result that completely matches those observed for touch (Crollen, Albouy, et al., 2017; Röder et al., 2004). Moreover, this outcome parallels other findings suggesting that somatosensory perceptual abilities of early blind people mostly rely, at least in similar tasks, on somatotopic reference frames (Collignon, Charbonneau, Lassonde, & Lepore, 2009; Crollen & Collignon, 2012; Roder, Focker, Hotting, & Spence, 2008). Therefore, it can be suggested that, despite the great relevance of the spatial representation of nociceptive painful stimuli in terms of behavioural adaptation, the ability to map nociceptive inputs according to spatiotopic reference frames is not innate but would rather be shaped by visual experience during development. Given the assumed similarities between the spatial representations of touch and nociception (Gallace, Torta, Moseley, & Iannetti, 2011; Legrain & Torta, 2015; Torta et al., 2013), the present data support the importance of other sensory modality inputs such as visual inputs for the development of the nociceptive system. Our results are indeed in line with studies that demonstrated a close interaction between nociceptive and visual stimuli occurring near the body (De Paepe, Crombez, & Legrain, 2015, 2017; De Paepe, Crombez, Spence, & Legrain, 2014; Filbrich, Alamia, Blandiaux, Burns, & Legrain, 2017; Filbrich, Halicka, Alamia, & Legrain, 2018). Furthermore, seeing the limb on which nociceptive stimuli are applied and the posture of the limb have been shown to affect the brain responses elicited by nociceptive stimuli, the evaluation of their intensity and the perception of pain (Gallace et al., 2011; Longo, Betti, Aglioti, & Haggard, 2009; Mancini, Longo, Kammers, & Haggard, 2011; Torta et al., 2013). It is worth noting that our data shed more light on the reasons underlying the impact of body posture on the perception of pain (Gallace et al., 2011; Torta et al., 2013). Indeed, it has been suggested that the decreased performance during somatosensory TOJ tasks with crossing hands reflects the cost of the automatic spatiotopic remapping, and the resources needed to solve the conflict between the different spatial representations in order to adequately localize the relevant stimulation (Shore et al., 2002). The influence of the posture on the subjective intensity might therefore reflects the additional cognitive load associated with the necessity to realign the spatiotopic map with the somatotopic one when the hands are crossed, resulting in limited neural resources available to process intensity feature (Torta et al., 2013). While further studies are needed to better characterize the relation between those two aspects, the afore-mentioned and the present data highlight the importance of the spatial representations of nociceptive inputs for the perception of pain. Being able to adequately defend the body against potential physical threats requires both coding somatosensory inputs and representing the body limbs according to their relative location in external space. This is requested for planning spatially-guided actions against those threats. In that sense, spatiotopic mapping of somatosensory inputs might be conceptualized as a premise of multisensory integration, facilitating interactions between somatic and non-somatic (e.g. visual) inputs. Spatiotopic mapping therefore offers a multimodal reference frame shared by the different sensory modalities. Nociceptive stimuli have been shown to interact with visual stimuli, especially with those occurring in the proximity of the limb on which the nociceptive stimuli are applied (De Paepe et al., 2015, 2017; De Paepe et al., 2014; Filbrich, Alamia, Blandiaux, et al., 2017; Filbrich et al., 2018). Accordingly, nociceptive stimuli interact with extra-somatic stimuli in the peripersonal reference frame, a representation integrating spatial information from the body space and spatial information from the external space immediately surrounding the body (Vallar & Maravita, 2009). Since the peripersonal reference frame is assumed to play an important role in shaping interactions between the body and external objects in contact with the body (Brozzoli et al., 2014), it is hypothesized that spatiotopic mapping of nociceptive stimuli represents an important process to integrate physical threats in peripersonal representations of the body, therefore optimizing defensive behaviours to protect the body against potential damages (Haggard et al., 2013; Legrain & Torta, 2015).

Based on the present data, we can hypothesize an involvement of premotor and posterior parietal areas in nociception and pain. These cortical regions have been indeed demonstrated to play a role in coding tactile inputs according to spatiotopic reference frames (Azanon, Longo, Soto-Faraco, & Haggard, 2010; Bolognini & Maravita, 2007; Crollen, Lazzouni, et al., 2017; Lloyd, Shore, Spence, & Calvert, 2003; Takahashi, Kansaku, Wada, Shibuya, & Kitazawa, 2013; Wada et al., 2012) and in participating in crossmodal interactions within peripersonal frames of reference (Avillac, Deneve, Olivier, Pouget, & Duhamel, 2005; Brozzoli, Gentile, Petkova, & Ehrsson, 2011; Graziano et al., 1997; Makin, Holmes, & Zohary, 2007). Premotor and posterior parietal areas have also been shown to be activated by nociceptive and painful stimuli (see review in Apkarian, Bushnell, Treede, & Zubieta, 2005). However, because their activity was interpreted as related to pain epiphenomena (Apkarian et al., 2005), they received much less attention than other activated cortical areas (Legrain, Iannetti, Plaghki, & Mouraux, 2011). For instance, a source modelling study of the magnetic fields elicited by nociceptive stimuli suggests a response in the posterior parietal cortex evoked in the same latency range than responses in primary and secondary somatosensory cortices (Nakata et al., 2008). Therefore, considering that the cortical areas classically observed in response to nociceptive and painful stimuli, such as cingulate and operculo-insular cortices, are often seen as belonging to a brain network involved in the detection of any sensory stimuli that might have an impact on the body’s integrity (e.g. Legrain et al., 2011), we might hypothesize that premotor and posterior parietal areas are associated to that network with the purpose of mapping physically threatening objects into external space and preparing spatially guided actions in order to protect the body.

The difference in representing spatially nociceptive inputs between early blind and normally sighted people illustrates the neuroplasticity of the nociceptive system and supports the idea that nociception might develop along different tracks, depending on sensory experience through the available sensory systems. This would lead to qualitatively different ways of processing threatening information and perceiving pain. Accordingly, several studies have assumed that early blind people might display differences in their sensitivity to pain as compared to normally sighted individuals (Slimani, Danti, Ptito, & Kupers, 2014; Slimani, Ptito, & Kupers, 2015). However, further reports suggested that hypersensitivity to pain in early blindness might be restricted to changes in the processing of C-fibre inputs (Slimani, Plaghki, Ptito, & Kupers, 2016) and would be related to differences in anxiety levels and attention between blind and sighted participants (Holten-Rossing, Slimani, Ptito, Danti, & Kupers, 2018). In the present study, we did not find any difference between the groups of participants regarding Aδ fibre detection thresholds, and blind and sighted participants perceived the nociceptive stimuli similarly. Nonetheless, further studies are needed to better characterize the influence of cross-modal plasticity of the nociceptive system on the perception of pain.

In conclusion, the present experiments emphasized the exteroceptive function of nociception by highlighting the importance of the spatial representations of nociceptive inputs and the role of vision for the development of the nociceptive system. The present study also indicates that visual deprivation can impact the organization of the cortical network underlying nociception, potentially leading to differences in pain processing in early blind people.

## 6. Acknowledgments

The authors thank A. Mouraux and A. Alamia (Institute of Neuroscience, Université catholique de Louvain) for their help in coding and writing the Matlab scripts related to the experiments.

## List of abbreviations

JND: just noticeable difference
SOA: stimulus onset asynchrony
PSS: point of subjective simultaneity
TOJ: temporal order judgement task

